# Propofol differentially modulates consolidation of schema-congruent and –incongruent memory

**DOI:** 10.1101/2025.03.13.643087

**Authors:** Lukas Risse, Deetje Iggena, Lili Landerer, Mario Menk, Heidi Olze, Daniel J. Salchow, Carsten Finke, Yee Lee Shing, Christoph J. Ploner

## Abstract

Congruency of newly learned information with previous knowledge (i.e. a mental schema) leads to facilitated encoding and rapid integration into neocortical memory networks. It is less known whether this is associated with a differential involvement of the hippocampus in consolidation of schema-congruent and –incongruent information. Here, we used the GABA_A_-ergic anesthetic propofol to transiently modulate hippocampal neural activity shortly after encoding of schema-congruent and -incongruent information in human patients. We found a significant difference in memory of schema-congruent and –incongruent words in patients that was absent in controls. This effect was driven by a benefit for schema-congruent words, thus suggesting that propofol administration facilitated consolidation of previously encoded schema-congruent items. Our results suggest that schema-congruency of newly learned information significantly modulates involvement of hippocampus-dependent networks during memory consolidation. They further support the hypothesis of a competitive interaction between hippocampus and extra-hippocampal networks during early memory consolidation.

## Introduction

Theories on systems memory consolidation converge on the idea that the passage of time alters how the brain represents conscious memories (Squire et al. 2015; Dudai et al., 2015; Barry & Maguire 2019; Moscovitch & Gilboa 2022). There is however an ongoing debate on the respective roles of the hippocampus and neocortex during systems consolidation. The heterogeneity of findings from studies of hippocampal-neocortical interaction suggests that memory consolidation is not a single process that stereotypically follows encoding but rather results from the interaction of distributed representations whose contributions are modulated by content-related and contextual factors (Dudai et al., 2015; Barry & Maguire 2019; Moscovitch & Gilboa 2022). One important factor may be the relatedness of new information to previous knowledge or mental schemas (Bartlett 1932; Tse et al., 2007). There is evidence from animal and human studies that these factors facilitate integration of new information into neocortical networks (van Kesteren et al. 2012; Hebscher et al., 2019; Alonso et al. 2020). In humans, EEG studies have shown that schema-congruency effects are already detectable at encoding of new information into memory (Packard et al. 2017). fMRI studies have further shown that availability of a schema significantly modulates connectivity between hippocampus and ventromedial prefrontal cortex at encoding and during the post-encoding period (van Kesteren et al. 2010; Liu et al. 2017; Audrain & McAndrews 2022; Guo et al. 2023). Neural activity in the hippocampus during this latter period may represent an important step in early memory consolidation as it predicts later memory performance across a variety of tasks and stimulus materials (Tambini & Davachi 2013; Tambini & D’Esposito 2020). So far, it is however unclear whether the hippocampus differentially contributes to consolidation of schema-congruent and –incongruent information (Hebscher et al., 2019).

Investigation of the role of the human hippocampus for early memory consolidation ideally requires techniques that modulate hippocampal neural activity without affecting prior encoding and later retrieval. However, the human hippocampus and surrounding structures are not directly accessible for current non-invasive brain stimulation techniques. Invasive deep brain stimulation of the human entorhinal cortex during memory tasks is limited to patients undergoing evaluation for epilepsy surgery and non-invasive electrical stimulation of the hippocampus is currently still in development (Titiz et al., 2017; Violante et al. 2023). An alternate approach to interfere transiently with hippocampal activity is the administration of the short-acting anesthetic propofol (2,6-diisopropylphenol). This drug acts as an agonist on the gamma-aminobutyric-acid (GABA)-A receptor and as a partial antagonist on N-methyl-D-aspartate (NMDA) receptors (Sahinovic et al., 2018; Walsh, 2018). PET and fMRI studies showed that propofol significantly modulates hippocampal metabolism and activation during memory tasks (Sun et al. 2008; Pryor et al., 2015). In two recent studies, human participants learned verbal or visuospatial material and received general anesthesia with propofol some minutes thereafter. After recovery from anesthesia, participants showed a memory pattern that suggested modulation of hippocampus-dependent early steps of memory consolidation (Moon et al., 2020; Iggena et al. 2022).

In the current project, we combined the propofol-approach established in previous work (Galarza Vallejo et al., 2019; Moon et al., 2020; Iggena et al. 2022) with a modification of a verbal schema memory task (Craik & Tulving 1975; Vogel et al. 2017; Packard et al. 2017). Neurologically normal patients undergoing short propofol anesthesia for minor surgery learned a list of randomized words that were either congruent with a soundscape provided by headphones or completely incongruent. About 12 minutes after learning, participants received anesthesia with intravenous propofol for about 60 minutes. Three hours after learning, participants were tested for recall and recognition of schema-congruent and –incongruent words. Performance was compared to participants receiving local anesthesia and to participants receiving no anesthesia. We reasoned that any differences in hippocampus-dependency of schema-congruent and -incongruent words during early consolidation should lead to differences in memory performance following anaesthesia.

## Results

### Learning

During the learning phase, word lists were presented repeatedly in three learning blocks with an equal number of randomized schema-congruent and -incongruent words (see Figure 1 for experimental protocol).

**Figure 1.**
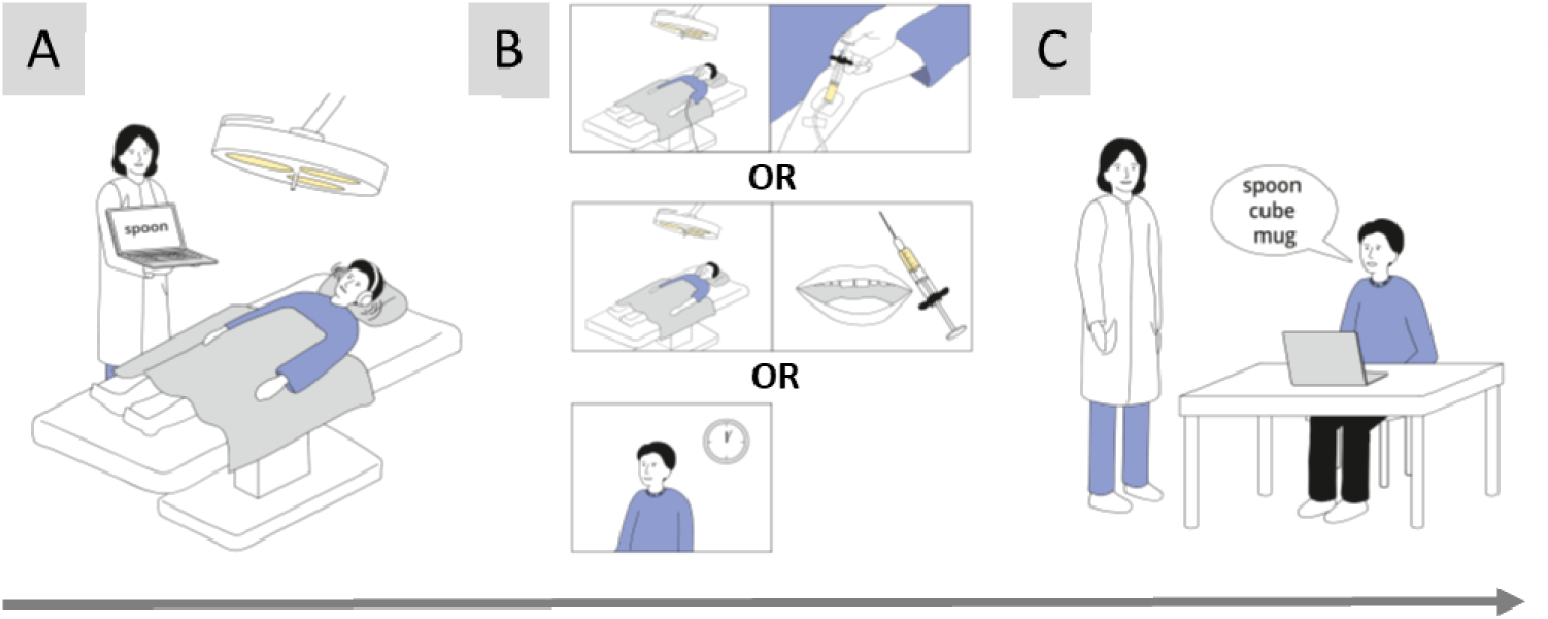
Task and experimental conditions. **A**, learning phase. While being in a supine position, participants learned a list of words that were either congruent with a predefined schema (“restaurant”) or not. Participants wore headphones that provided a soundscape suggestive of a restaurant. **B**, consolidation phase. About 12 minutes after learning, participants received general anesthesia with propofol for about 60 minutes (top row), local anesthesia (middle row) or no anesthesia (bottom row). **C**, retrieval phase. About three hours after learning, participants were tested for recall and recognition of words

All groups showed learning of word lists across the three learning blocks with a continuous increase of recalled words (Χ^2^(2) ≥ 36.4, p < 0.001, Friedman ANOVA, table 1). The increase from block 1 to block 3 was significant in all groups (Z ≤ - 1.72, p < 0.001, Wilcoxon signed ranks test). However, the percentage of learned words differed between groups within learning blocks and was significantly different in blocks 1 and 3 (block 1: Χ^2^(2) = 9.06, p = 0.011; block 2: Χ^2^(2) = 5.06, p = 0.08; block 3: Χ^2^(2) = 8.47, p = 0.014, Kruskal-Wallis ANOVA). Post-hoc testing showed that these differences were mainly due to differences between the propofol group and the no anesthesia group (block 1: Z = −2.95, p = 0.003; block 3: Z = −2.65, p = 0.008; Mann-Whitney test). Performance between the propofol group and the local anesthesia group did not differ significantly (block 1: Z = −1.29, p = 0.2; block 3: Z = - 2.95 p = 0.28; Mann-Whitney test). We suppose that presurgical arousal may have affected performance in patients.

**Table 1.** Performance of subject groups during learning blocks in percent correct. Values are medians and interquartile ranges. p-values refer to Friedman-ANOVA.

|  | Block 1 | Block 2 | Block 3 | p |
| --- | --- | --- | --- | --- |
| Propofol | 22.5 (20.0 – 32.5) | 42.5 (27.5 – 47.5) | 42.5 (30.0 – 62.5) | <0.001 |
| No anesthesia | 32.5 (25.0 – 42.5) | 50.0 (32.5 - 72.5) | 62.5 (55.0 – 75.0) | <0.001 |
| Local anesthesia | 27.5 (22.5 – 35.0) | 40.0 (35.0 – 52.5) | 52.5 (42.5 – 62.5) | <0.001 |

When analyzed separately, schema-congruent words were significantly better learned than schema-incongruent words in all groups (performance averaged across blocks; propofol group: Z = - 3.55, p < 0.001; no anesthesia group: Z = - 2.26, p = 0.024; local anesthesia group: Z = - 3.72, p < 0.001; Wilcoxon signed ranks test; figure 2). Since schema-congruent and -incongruent words were matched in terms of word length, emotional valence, arousal, imageability and frequency of occurrence in German language (see supplement), these differences suggest that the contextual factors of our paradigm reliably activated knowledge from previous experience that supported learning of new schema-congruent information.

**Figure 2.**
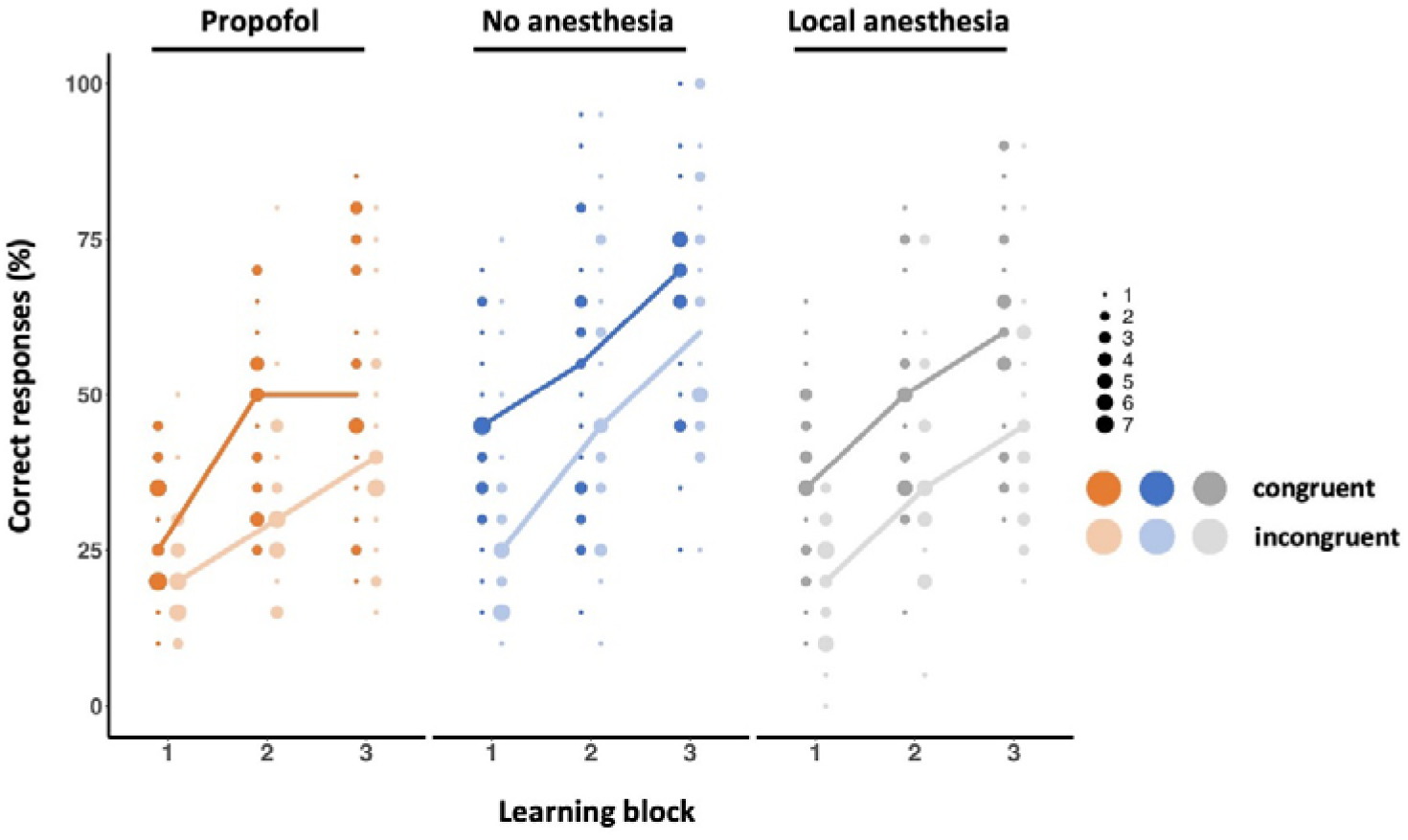
Performance of subject groups during learning blocks in percent correct responses. Performance separately for schema-congruent words (dark dots) and incongruent words (light dots). Dot size shows number of identical values. Lines connect median values.

### Recall

All subject groups recalled at least 30% of the learned words across the three-hour delay between the end of learning (i.e. block 3) and recall testing, regardless of schema congruency (table 2).

**Table 2.**
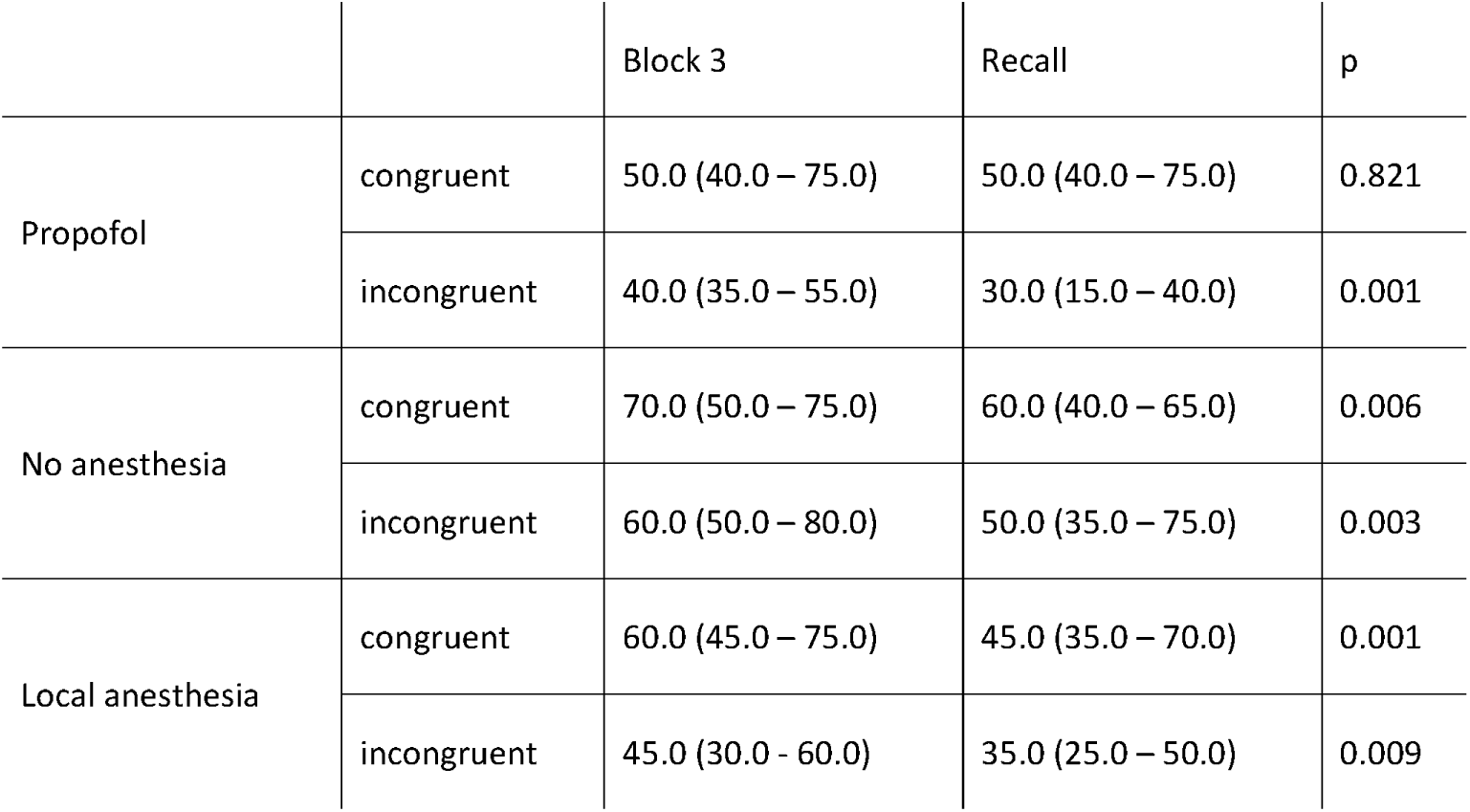
Performance of subject groups in the last learning block (block 3) and at delayed recall about 3 hours later in percent correct. Values are medians and interquartile ranges. p-values refer to Wilcoxon signed ranks test.

|  |  | Block 3 | Recall | p |
| --- | --- | --- | --- | --- |
| Propofol | congruent | 50.0 (40.0 – 75.0) | 50.0 (40.0 – 75.0) | 0.821 |
|  | incongruent | 40.0 (35.0 – 55.0) | 30.0 (15.0 – 40.0) | 0.001 |
| No anesthesia | congruent | 70.0 (50.0 – 75.0) | 60.0 (40.0 – 65.0) | 0.006 |
|  | incongruent | 60.0 (50.0 – 80.0) | 50.0 (35.0 – 75.0) | 0.003 |
| Local anesthesia | congruent | 60.0 (45.0 – 75.0) | 45.0 (35.0 – 70.0) | 0.001 |
|  | incongruent | 45.0 (30.0 – 60.0) | 35.0 (25.0 – 50.0) | 0.009 |

However, significant differences were found between performance on block 3 and at recall testing for schema-incongruent words in all groups (Z ≤ −2.6, p ≤ 0.009, Wilcoxon signed ranks test). Similarly, performance on block 3 and at recall testing differed for schema-congruent words in the no anesthesia group and the local anesthesia group (Z ≤ −2.74, p ≤ 0.006, Wilcoxon signed ranks test), but not in the propofol group (Z = −0.23, p = 0.821, Wilcoxon signed ranks test).

We then analyzed possible differences in forgetting of schema-congruent and - incongruent words across the three-hour delay. We subtracted block 3 performance values from recall values, thus yielding negative Δ values for forgetting (figure 3). In the no anesthesia group and the local anesthesia group, schema-congruent and – incongruent words were equally forgotten with no significant difference between Δ values (figure 3, no anesthesia group: Z = −0.53, p = 0.6; local anesthesia group: Z = 0.19, p = 0.85, Wilcoxon signed ranks test). By contrast, in the propofol group, a significant difference in forgetting between the two stimulus categories was found (Z = −3.27, p = 0.001, Wilcoxon signed ranks test) with forgetting of schema-incongruent words and almost unchanged performance for schema-congruent words, despite general anesthesia (figure 3).

**Figure 3.**
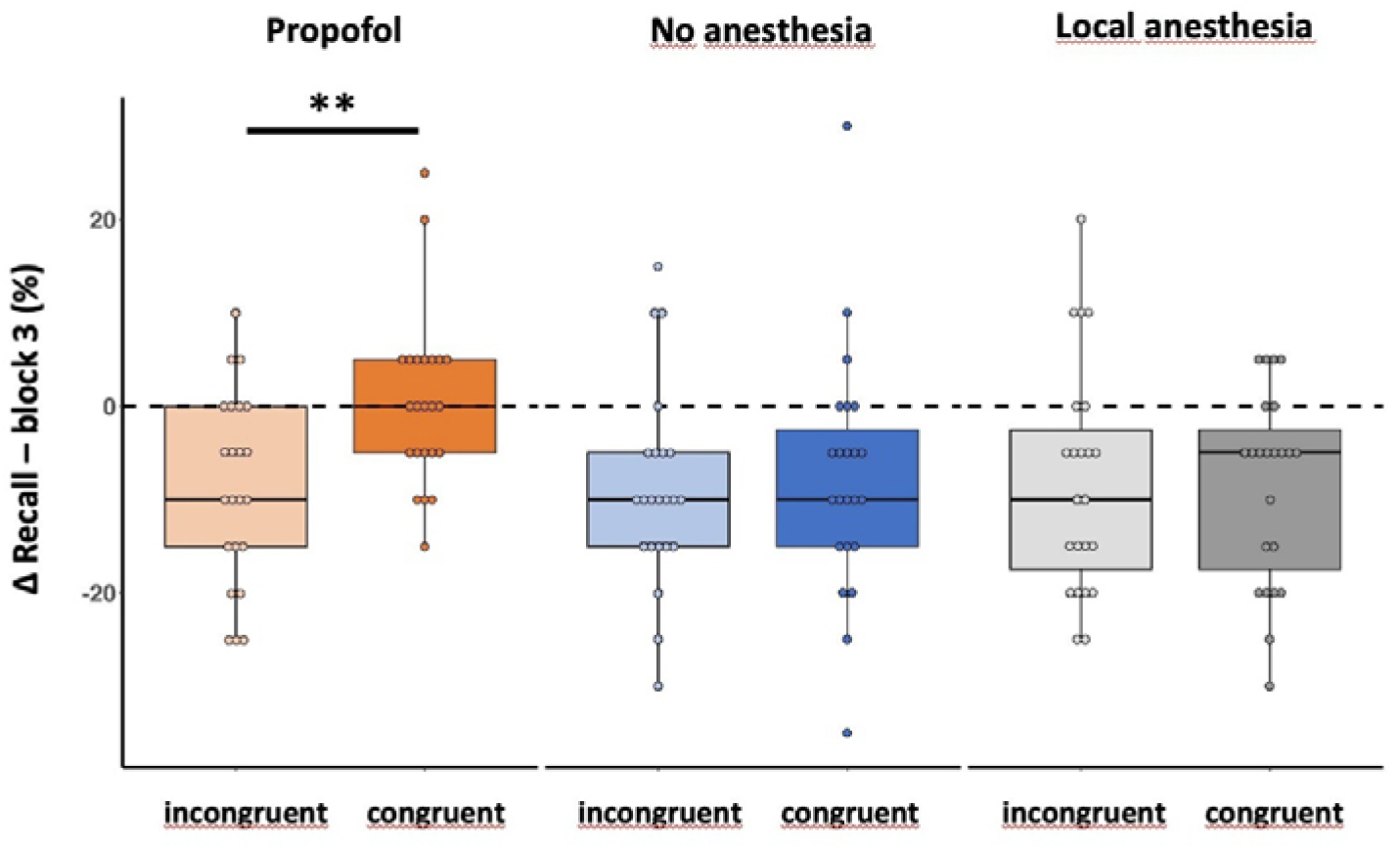
Differences in recall of stimulus words from learning block 3 to recall three hours after learning. separately for schema-congruent and -incongruent words for each group. Negative values denote forgetting. Dashed line indicates unchanged performance. ** p = 0.001, Wilcoxon signed ranks test

Furthermore, while Δ values for incongruent words did not differ significantly between groups, Δ values for congruent words differed significantly (incongruent: Χ^2^(2) = 0.70, p = 0.966; congruent: Χ^2^(2) = 9.47, p = 0.009, Kruskal-Wallis ANOVA). Post-hoc testing showed that Δ values differed significantly between the propofol group and the other two groups (propofol – no anesthesia: Z = −2.72, p = 0.007; propofol – local anesthesia: Z = −2.60, p = 0.009; Mann-Whitney test). No difference was found between the no anesthesia and the local anesthesia group (Z = −0.12, p = 0.902; Mann-Whitney test).

## 3. Recognition

Like in a previous study, we found no effect of propofol on recognition memory of word lists (Moon et al. 2020). Recognition performance and confidence ratings were almost at ceiling in all subject groups, both for schema-congruent and -incongruent words (table 3). In addition, no significant differences were found between the two stimulus categories in all subject groups (recognition: Z ≥ −1.65, p ≥ 0.098; familiarity: Z ≥ −0.84, p ≥ 0.398, Wilcoxon signed ranks test). In addition, although the absolute number of false positive ratings was low, we found significantly more false positive ratings for schema-congruent words in all subject groups (table 3; Z ≤ −2.41, p ≤ 0.016, Wilcoxon signed ranks test), thus suggesting that schema-dependent predictive processing was still implicitly present in the testing period of our paradigm (Hubbard et al. 2019; Höltje & Mecklinger 2022).

**Table 3.**
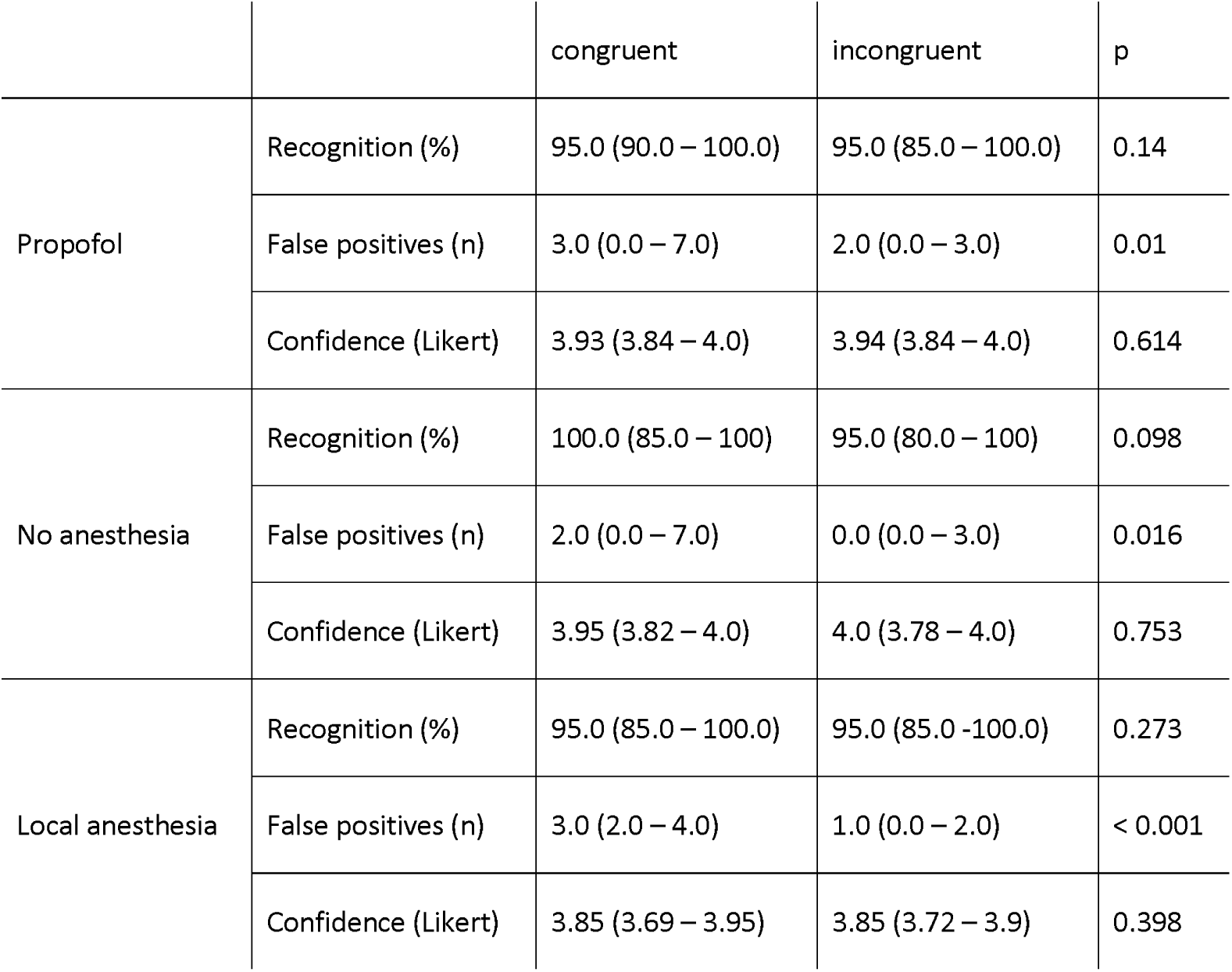
Performance of subject groups at recognition testing. Values are medians and interquartile ranges. p-values refer to Wilcoxon signed ranks test

|  |  | congruent | incongruent | p |
| --- | --- | --- | --- | --- |
| Propofol | Recognition (%) | 95.0 (90.0 – 100.0) | 95.0 (85.0 – 100.0) | 0.14 |
|  | False positives (n) | 3.0 (0.0 – 7.0) | 2.0 (0.0 – 3.0) | 0.01 |
|  | Confidence (Likert) | 3.93 (3.84 – 4.0) | 3.94 (3.84 – 4.0) | 0.614 |
| No anesthesia | Recognition (%) | 100.0 (85.0 – 100) | 95.0 (80.0 – 100) | 0.098 |
|  | False positives (n) | 2.0 (0.0 – 7.0) | 0.0 (0.0 – 3.0) | 0.016 |
|  | Confidence (Likert) | 3.95 (3.82 – 4.0) | 4.0 (3.78 – 4.0) | 0.753 |
| Local anesthesia | Recognition (%) | 95.0 (85.0 – 100.0) | 95.0 (85.0 -100.0) | 0.273 |
|  | False positives (n) | 3.0 (2.0 – 4.0) | 1.0 (0.0 – 2.0) | < 0.001 |
|  | Confidence (Likert) | 3.85 (3.69 – 3.95) | 3.85 (3.72 – 3.9) | 0.398 |

## Discussion

The present study investigated effects of the anaesthetic propofol on memory consolidation of to-be-remembered words that differ in their congruency with a predefined schema. For post-anesthesia recall, we found a significant difference between schema-congruent and schema-incongruent words. This was mainly due to a benefit for schema-congruent words uniquely in the propofol group, thus suggesting that propofol facilitated consolidation of newly encoded schema-congruent words but not of schema-incongruent words. Our findings suggest that schema-congruency modulates involvement of hippocampus-dependent networks during early memory consolidation. They are further consistent with a competitive interaction of two memory systems, with an inhibitory role of hippocampus-dependent networks on extrahippocampal regions in the post-encoding period.

Although not explicitly instructed, the results of the learning trials show that the contextual factors of our paradigm (i.e. the soundscape and the large proportion of stimulus words consistent with a single context) reliably activated a mental schema that supported learning of new information. Consistent with the hypothesis that schemas provide a framework that facilitates encoding (van Kesteren et al. 2012; Hebscher et al. 2019; Sekeres et al. 2024), a significant benefit for schema-congruent words across learning blocks was observed in all three groups. It is unclear whether this mainly reflects a quantitative difference in processing of schema-related and -unrelated information or whether it also reflects distinct trajectories of consolidation recruiting distinct neural substrates. fMRI studies have shown that availability of a schema significantly modulates connectivity between hippocampus and ventromedial prefrontal cortex at encoding and during the post-encoding period (van Kesteren et al. 2010; Liu et al. 2017; Audrain & McAndrews 2022; Guo et al. 2023). There is however no clear behavioral evidence from humans whether and how schema-congruency alters the role of the hippocampus during consolidation. In particular, it is open whether rapid integration of schema-congruent information into neocortical networks also means a change in the temporal properties of hippocampal involvement during consolidation (Hebscher et al. 2019). If so, behavioral consequences of modulation of hippocampal neural activity during the consolidation period should be different for schema-congruent and -incongruent information. The pharmacological manipulation used here used a potent anesthetic that targets α5GABA_A_-receptors, i.e. inhibitory receptors that are particularly abundant in the hippocampus (Engin et al. 2018; Sperk et al. 2020). Accordingly, propofol administration has been shown to modulate key processes of memory formation in animal experiments (Wei et al. 2002; Nagashima et al. 2005; Takamatsu et al. 2005), hippocampal activity in human fMRI studies (Pryor et. 2015) and hippocampus-dependent memory consolidation in humans (Moon et al. 2020; Iggena et al. 2022). The observed differences in free recall of schema-congruent and - incongruent words after propofol administration thus are consistent with the hypothesis that schema-dependent differences in memory consolidation are not a mere consequence of facilitated encoding of schema-congruent information but rather result from a continuous process that significantly extends into the post-encoding period and is accompanied by a different role of the hippocampus for consolidation of schema-congruent and -incongruent words.

The unimpaired retention of schema-unrelated words in the propofol group seems to contradict earlier results from a previous study, where application of propofol impaired free recall of word lists learned shortly before injection (Moon et al. 2020). However, although both studies used word lists as stimulus material, the mode of presentation of the lists was significantly different and may have affected how stimuli were encoded and consolidated. In our first study, stimuli consisted of a simple list of words from various semantic categories that was repeatedly presented in a fixed order during learning (Moon et al. 2020). In the present study, lists consisted of words half of which were congruent with a pre-defined schema that were presented pseudo-randomly in unpredictable order during learning. Since the experiments of Donald Hebb it is known that serial learning benefits significantly from repeated presentation of stimulus material in a fixed order (Hebb, 1961). This effect has been observed across a wide range of verbal and visual stimuli and various associative mechanisms have been proposed to account for it (Araya et al., 2024). In principle, individual items may be associated with a distinct position in the list or with neighbouring items. Serial recall experiments suggest that it is more likely that items of a list may be grouped into several chunks of information or into a unified representation of the entire list (Burgess & Hitch, 2006; Page & Norris, 2009). These associative mechanisms are not necessarily mutually exclusive and may have contributed to performance in our first study (Moon et al., 2020) but are unlikely in the present study. Experiments with transcranial brain stimulation following visual paired associate learning and propofol injections following navigational learning suggest that the formation of associations between memory items critically depends on neural activity within a time window of about 60 Minutes after learning (Tambini & D’Esposito, 2020; Iggena et al., 2022). Therefore, one explanation for the seemingly divergent findings from our first study and the present investigation is that propofol may not have interfered with memory of single items in the first study but rather with associative mechanisms that support memory consolidation in conditions where the stimulus material is learned repeatedly in a fixed order. In support of this hypothesis, studies have shown that consolidation of spontaneous associative binding of words in repeated word list learning is impaired in patients with lesions of the hippocampus (Grewe et al., 2020). Accordingly, recall performance levels following propofol anaesthesia in our first study (Moon et al., 2020) are similar to recall before and after propofol anaesthesia in the present study.

So why then did we observe better memory for schema-congruent words rather than impaired memory for schema-incongruent words after administration of an inhibitory drug? Human studies have shown that various drugs, including alcohol and benzodiazepines, can lead to better verbal and visual memory performance when administered shortly after learning (e.g. Parker et al. 1981; Ghoneim et al. 1984; Fillmore et al. 2001; Carlyle et al. 2017). One common denominator behind these effects is suggested to be a GABA_A_-mediated mechanism, whereby these drugs inhibit induction of long-term potentiation (LTP) in the hippocampus (Blitzer et al. 1990; del Cerro et al. 1992). The resulting inhibition of hippocampal encoding would prevent retroactive interference that would otherwise have weakened memories established prior to drug administration (Mednick et al. 2011). In line with this hypothesis, a recent study reported effects of intrahippocampal injections of the GABA_A_-agonist muscimol on performance of rats in a novel-object recognition memory task (Sawangjit et al. 2022). About 30 minutes after encoding, rats received muscimol injections either in the awake state or during sleep. At testing, opposed effects were observed with decreased memory for objects when muscimol was applied during sleep and increased memory when it was applied during the awake state. The authors hypothesized a competitive interaction between a hippocampally-mediated memory system and memory systems in extrahippocampal brain regions. In the awake state, consolidation processes in these latter regions might be disturbed by interfering hippocampal activity (Sawangjit et al. 2022). Collectively, these and the aforementioned findings are consistent with an “opportunistic” hypothesis of memory consolidation, which posits that both synaptic and systems consolidation of newly encoded information are facilitated by subsequent periods of reduced interference – be it by slow-wave sleep or by application of drugs (Mednick et al. 2011).

When our results are discussed within the framework of opportunistic memory consolidation, propofol-induced decreased forgetting of schema-congruent words would mean that – despite presumed facilitation of neocortical integration – hippocampal networks are still relevant for consolidation of schema-congruent words in the post-encoding period. This role seems to be inhibitory, at least at this timepoint and in our paradigm. As a consequence, hippocampal deactivation may lead to facilitated extrahippocampal integration of schema-congruent words, while schema-incongruent words do not benefit. Our findings may thus support the hypothesis that hippocampus-dependent and extrahippocampal memory networks can compete in the post-encoding period (Poldrack & Packard 2003; Sawangjit et al. 2022). How this relates to changes in connectivity between hippocampus and neocortex is not clear, as opposing connectivity patterns between hippocampus and ventromedial prefrontal cortex have been reported to be associated with memory of schema-congruent information in the post-encoding period (van Kesteren et al. 2010; Bein et al. 2014; Liu et al. 2017; Audrain & McAndrews 2022; Guo et al. 2023).

In conclusion, our results show that schema-congruency affects early consolidation of verbal information. The pattern of results is consistent with a differential role of hippocampus-dependent memory networks for schema-congruent and –incongruent information shortly after learning. Our data further suggest that despite accelerated encoding, integration of schema-congruent words into presumably neocortical networks seems to significantly extend into the post-encoding period, with an inhibitory role of hippocampus-dependent memory networks. These results thus add evidence to the concept of competing hippocampal and extra-hippocampal memory systems that flexibly interact according to availability of previous knowledge.

### Limitations of the study

Our neuro-pharmacological approach only allows for indirect inferences about the brain systems involved. Conclusions should therefore be substantiated by additional imaging experiments. Furthermore, propofol-induced decreased forgetting of schema-congruent words needs not result from distinct processing of schema-congruent and -incongruent words. Both stimulus categories may be processed and represented by the same brain systems but may show different levels of semantic interference, with a significant reduction in interference by propofol for schema-congruent words only. Previous research indeed has shown that semantically related verbal and visual information may interfere in memory, even when presented with unrelated intervening trials (Harvey & Schnur 2016; Wei & Schnur 2016). However, if this factor would have been decisive, we would have expected superior learning and memory of schema-incongruent words. We therefore favour the hypothesis of distinct processing modes for both stimulus categories.

## Resource availability

### Lead contact

Requests for further information and resources should be directed to and will be fulfilled by the lead contact, Christoph Ploner.

### Materials availability

Experimental materials consisted of a word list that is provided in the supplemental information of the manuscript.

### Data and code availability

De-identified human data have been deposited to an OSF repository and are publicly available. The DOI is listed in the key resources table. This paper does not report original code. Any additional information required to reanalyze the data reported in this paper is available from the lead contact upon request.

## Acknowledgments

This study was funded by the Deutsche Forschungsgemeinschaft (DFG, German Research Foundation), Project number 327654276 - B05, SFB 1315. We thank the study participants and the staff of the Department of Otorhinolaryngology, the Department of Ophthalmology and the Department of Anesthesiology of the Charité – Universitätsmedizin Berlin, Campus Virchow-Klinikum, and Stephan Kinder, Troisdorf, for their kind support of our study. Miriam Seith assisted during preparation of figures. Special thanks to Lynn Nadel for helpful comments on the manuscript.

## Author contributions

Conceptualization, C.J.P., L.R., and Y.L.S; Methodology, C.J.P., L.R., D.I., M.M., C.F., Y.L.S.; Investigation, L.R. and L.L.; Writing – Original Draft, C.J.P. and L.R.; Writing – Review & Editing, C.J.P., L.R., Y.L.S., D.I., and C.F.; Funding Acquisition, C.J.P. and C.F.; Resources, M.M., H.O., and D.J.S.; Supervision, C.J.P.

## Supplemental information

Document S1

## KEY RESOURCES TABLE

| REAGENT or RESOURCE | SOURCE | IDENTIFIER |
| --- | --- | --- |
| Deposited data |  |  |
| Experimental data | This paper | https://osf.io/gv9zs/ |
| Software and algorithms |  |  |
| IBM SPSS Statistics for Windows, Version 29.0.2.0 | Armonk, NY: IBM Corp | https://www.ibm.com |
| Presentation | Neurobehavioral Systems, Berkeley, CA | https://www.neurobs.com/ |

### EXPERIMENTAL MODEL AND STUDY PARTICIPANT DETAILS

A total of 69 participants between 18 and 60 years without any history of neurological or psychiatric disorders, hearing disorders, visual disturbances or substance abuse was included in the study. All participants were native German speakers. Three groups of 23 participants each, matched for sex, age and educational level were tested with a verbal memory task (table 4). One group received general anesthesia with propofol between learning and testing, one group local anesthesia and one group no anesthesia.

**Table 4.** Demographic and clinical data of the invested patient groups. Values are medians and interquartile ranges; n.a. not applicable.

|  | Propofol | No anesthesia | Local Anesthesia |
| --- | --- | --- | --- |
| n | 23 | 23 | 23 |
| Female / Male | 10 / 13 | 13 / 10 | 13 / 10 |
| Age (Years) | 37 (21.5-49.5) | 39 (25.5-52.5) | 41 (29-51) |
| Years of Education | 15 (14-17) | 16 (15-18) | 15 (14-16.5) |
| Medical Procedure | Strabismus surgery<br>(n =19)<br>Maxillofacial surgery<br>(n=4) | n.a. | Maxillofacial surgery<br>(n=23) |
| Propofol Bolus Dose<br>(mg) | 200 (200-200) | n.a. | n.a. |
| Propofol Maintenance<br>Dose (mg/kg/h) | 6 (6-6) | n.a. | n.a. |
| Remifentanil Dose<br>(µg/kg/min) | 0.2 (0.2-0.2) | n.a. | n.a. |
| Time between end of<br>learning and Propofol<br>(min) | 12 (9-14.5) | n.a. | n.a. |
| Duration anesthesia<br>(min) | 61 (56-66.5) | n.a. | n.a. |
| Time between end of<br>anesthesia and testing<br>(min) | 124 (120.5-141) | n.a. | n.a. |
| Time between end of<br>learning and testing<br>(min) | 195 (185.5-202) | 189 (176-199) | 192 (184-202) |

The propofol group consisted of participants undergoing general anesthesia with propofol for minor strabismus surgery or minor maxillofacial surgery, such as nasal septum reposition and material removal (table 4). All patients received the same anesthetic protocol. The no anesthesia group consisted of participants that underwent no surgical or other medical procedures (table 4). The local anesthesia group consisted of participants undergoing local anesthesia for minor maxillofacial surgery, such as wisdom tooth resection or dent implantation (table 4). This group was recruited to control for possible pre-surgical arousal effects on memory task performance (Pryor et al, 2015).

Participants undergoing anesthesia were recruited during preparatory outpatient visits. The no anesthesia group was recruited via the intranet of the Charité - Universitätsmedizin Berlin. Sample size was estimated prior to analysis based on data from previous studies on propofol effects on memory (Moon et al., 2020; Iggena et al. 2022). All procedures were approved by the ethics committee of the Charité – Universitätsmedizin Berlin. Every participant gave written consent before participation.

## METHOD DETAILS

### Behavioral testing

Participants were informed that they should perform a memory task and that they would receive a short additional test three hours later. Participants were not instructed about the purpose of the task and received no information about the semantic categories of the to-be-remembered words. Stimulus words were either congruent with the semantic category “restaurant” or from completely unrelated semantic categories. Words were presented visually on a 14-inch notebook computer at a distance of about 60 cm from the subject’s eyes, while participants were in a supine position. Words were composed of white letters against a black background. During the entire learning phase, participants wore noise-cancelling on-ear-headphones and were exposed to a soundscape suggestive of a restaurant. The soundscape was provided to facilitate the use of the schema “restaurant” during encoding of the words. Stimuli were programmed using Presentation software (Neurobehavioral Systems, Berkeley, CA, USA).

Stimuli consisted of 40 German words taken from the “Berlin Affective Word List Reloaded” (BAWL-R, Vo et al., 2009). Twenty words belonged to the semantic category “restaurant” (“restaurant words”, e.g. sauce, spice, fork) and 20 words to distinct and semantically unrelated categories (“non-restaurant words”, e.g. thumb, shower, magnet). Restaurant and non-restaurant words were matched for frequency of occurrence in German language, visual imaginability, arousal and emotional valence (see supplement for stimuli and matching procedure).

Before learning, the task was explained with written instructions on the notebook screen. Participants were then allowed to ask questions, until the examiner felt confident about participants’ comprehension of the task. Stimuli were presented in three blocks of 40 trials. Each restaurant and non-restaurant word was presented once in each block. Trial order was varied pseudo-randomly between blocks. During each trial, words were presented for 2000 ms, followed by a fixation cross for 2000 ms. After each block, participants were tested for learning of the words and were asked to freely recall as many words as possible. Responses were recorded for later offline analysis.

After learning, participants underwent general anesthesia (propofol group), local anesthesia (local anesthesia group) or were free to ambulate in the hospital (no anesthesia group). About three hours after the end of the learning phase, all participants were tested for recall and recognition of the learned words. Testing was conducted with the notebook in a quiet room with participants being in a seated position. No soundscape was presented. Initially, participants were requested to recall the word list and to report all recalled words orally while responses were recorded. For recognition, participants were presented a list of 80 words. The list consisted of the original list of 40 words, 20 new restaurant-related words and 20 new non-restaurant-related words in pseudorandom order. Words were presented successively on the notebook screen and participants were requested to decide by keypress whether a word had been part of the initial list or not. Presentation of a word was terminated by the keypress of the participant. In addition, participants were requested to rate the confidence of their decisions on a Likert scale from one (not confident) to four (absolutely confident). Then, a fixation cross was presented for 2000 ms and the next word was presented.

### Procedure

In the propofol group, participants performed the learning phase of the task in a preparation room or corridor adjacent to the operating theatre, while being in a supine position. The notebook screen was positioned over the participants’ head to ensure unrestricted and comfortable reading of the stimulus words. After termination of the learning phase, preparation for anesthesia started, participants received a peripheral venous access and had a final check-up talk with the responsible anesthesiologist. Participants were then transferred to the surgical theatre and anesthesia was induced with a bolus of 150 - 250 mg propofol, adjusted to the patient’s weight, followed by a continuous infusion of 6mg/kg/h propofol and 0,2 µg/kg/min remifentanil. The median time between the end of the learning phase and the injection of propofol was 12 minutes (IQR 9 – 14.5, table 4). During anesthesia, participants underwent surgery and were mechanically ventilated with a laryngeal mask. Median duration of anesthesia was 61 minutes (IQR 56 – 66.5, table 4). After surgery, participants were transferred to a recovery room where they were observed for about one hour. Post-surgery pain was treated with Paracetamol and Ibuprofen. Finally, participants were transferred to the ward, where they were tested for recognition and recall.

In the no anesthesia group, participants performed the learning, recall and recognition phases of the task in a doctor’s room equipped with an examination couch. Participants were put in a supine position. The notebook screen was positioned over the participants’ head to ensure unrestricted and comfortable reading of the stimulus words. After termination of the learning phase, participants were free to ambulate in the hospital, but were told to come back in 170 minutes to perform the final parts of the task. During this period, participants were not allowed to consume caffeine, drugs or other centrally acting substances.

In the local anesthesia group, participants performed the learning phase of the experiment in the surgical theatre after being prepared for surgery and while being in a supine position. The notebook screen was positioned over the participants’ head to ensure unrestricted and comfortable reading of the stimulus words. After the end of the learning phase, the surgeon and his team entered the room and participants had a final check-up talk. Then, local anesthesia was started with local injections of articain. The median time between the end of the learning phase and local anesthesia injection was 10 minutes (IQR 8 - 12). Post-surgical pain was treated with Ibuprofen. After surgery, participants either waited in a patient lounge or ambulated freely. Participants were told to come back 170 minutes after the end of the learning phase. Similar to the no anesthesia group, the free recall and recognition phases of the task were performed in a doctor’s room equipped with an examination couch.

## QUANTIFICATION AND STATISTICAL ANALYSIS

Data were analyzed by using IBM SPSS statistics (version 29.0). Memory performance was described as percent correct responses in each subject. For learning and delayed recall, we analyzed the percentage of correctly recalled items from the word list. For delayed recognition, we analyzed the percentage of correctly recognized words and the number of false positive recognitions for each subject. Group averages are given as medians with interquartile ranges. Since Kolmogorov– Smirnov testing showed that the assumption of a normal distribution had to be rejected for most variables, non-parametric statistical testing was used for statistical analysis (Altman, 1991; Altman and Bland, 2009). For analysis of within-group differences, we used Friedman-ANOVA and two-tailed Wilcoxon signed ranks tests. For analysis of between-group differences, we used Kruskal–Wallis ANOVA and two-tailed Mann–Whitney tests. Significance was accepted at a p < 0.05 level.

