## Supplemental information for "Propofol differentially modulates consolidation of schema-congruent and –incongruent memory"

**Stimulus selection**

*Rationale*

To study schema effects on early consolidation of memory, we used a word list paradigm that builds on previous work of schema effects on verbal memory (Craik & Tulving 1975; Vogel et al. 2017; Packard et al. 2017). In order to relate possible differences between memory of schema-congruent and –incongruent words unequivocally to their schema-congruency, we matched stimulus words from these two word categories as closely as possible in terms of number of syllables, word length, emotional valence, arousal, imageability and frequency of occurrence in German language. To this end, we based matching of words across stimulus categories primarily on The Berlin Affective Wordlist-Reloaded (BAWL-R), i.e. a list of German words that provides these variables (Kuchinke et al., 2006; Võ et al., 2009). However, since not all stimulus words were included in the BAWL-R, we conducted an additional online survey to assess emotional valence, arousal and imageability of the remaining stimulus words (see below). To match for frequency of occurrence in German language, we used the Korpusbasierte Wortgrundformenliste (DeReWo), a list of German words that provides word frequencies in German language (Leibniz-Institut für Deutsche Sprache, 2013).

*Selection of schema*

In order to compare memory of schema-congruent and –incongruent words, a schema with high imageability and familiarity across participants from various social and cultural backgrounds was needed. We deliberately avoided schema that may relate to individual negative experiences (e.g. school, hospital) or to distinct socio-biographical backgrounds (e.g. farm, forest). To ensure that the schema “restaurant” met criteria for our study, a survey with 22 participants was conducted on nurses, doctors, administrative staff and hospital visitors at the Charité – Universitätsmedizin Berlin (mean age 34 years, SD 11.6). Participants were asked to name the first words that come to their mind when thinking of a restaurant. A total of 183 different words was named by participants (mean: 22 words/participant, SD 4.8). Participants were further asked to rate familiarity and imageability of the term “restaurant” and whether they rate it as positive or negative. All participants stated to be familiar with and having a vivid imagination of a restaurant environment. All but one participant rated the term “restaurant” positively. Due to its high imageability and familiarity as well as its positive emotional valence, we decided to use the schema “restaurant” for our study.

*Selection of schema-congruent stimulus words*

Out of the total of 183 restaurant-related words named by participants, we selected the 50 most frequently mentioned German nouns with up to two syllables. We then searched the BAWL-R to determine their imageability, emotional valence, arousal, number of letters, word type and frequency of occurrence in German language. Thirty out of 50 words were found. The remaining 20 words were rated in an online survey with 30 randomly selected users of the online platform Prolific (<https://www.prolific.com/>) (mean age 35 years, SD 10.0). The survey was similar to the BAWL-R, but focused on emotional valence, arousal and imageability of these words. To assess the comparability between this additional survey and the survey conducted for the BAWL-R, 15 non-schema words that had already been rated in the BAWL-R were added to the new survey. The mean deviation between the 15 words already rated in the BAWL-R compared to their new rating in the online survey was 7,4% (SD 6,7) for emotional valence, 8,4% (SD 4,7) for arousal and 10,4% (SD 6,6) for imageability. The new online survey was thus considered comparable to the BAWL-R.

*Selection of schema-incongruent stimulus words*

We then searched for 50 German nouns with up to two syllables that matched the 50 previously defined restaurant-related words in terms of emotional valence, arousal, imageability, and frequency of occurrence in German language. We therefore searched the BAWL-R and DeReWo databases to find the non-restaurant word with the least deviation for the given variables from each restaurant word. This procedure resulted in a list of word-pairs, each consisting of a restaurant-related word and a matched non-restaurant-related word.

The 50 word-pairs were then rated regarding their relatedness to the schema “restaurant” on a Likert scale from 1 (no relation to restaurant) to 5 (strong relation to restaurant). The ratings were carried out on Prolific with 30 randomly selected test subjects (15 female, mean age 32 years, SD 8.5). The 40 word-pairs with the most strongly restaurant-related and least restaurant-related words were then used for the final stimulus set (mean ratings restaurant words = 4.75, SD = 0.25; mean ratings non-restaurant words = 1.44, SD = 0.29; p < 0.0001).

The final set of words showed no significant differences regarding emotional valence (mean difference on a scale from -3 to 3: 0.18, SD = 0.23; p = 0.95), arousal (mean difference on a scale from 1 to 5: 0.13, SD = 0,13 , p = 0.89), imageability (mean difference on a scale from 1 to 7: 0.2, SD = 0.16, p = 0.7) and frequency of occurrence in German language (mean difference on scale from 0 to 29: 1.23, SD = 1.27, p = 0.6).

| **Restaurant-related words** | **Non-restaurant-related words** |
| --- | --- |
| BAR | BACH |
| BIER | BAHRE |
| BROT | BETT |
| DURST | BLÜTE |
| ESSEN | BRIEF |
| FLASCHE | BRÜCKE |
| GABEL | BRUNNEN |
| GAST | BÜRSTE |
| GERUCH | DAUMEN |
| GEWÜRZ | DUSCHE |
| GLAS | ENKEL |
| HERD | FICHTE |
| HUNGER | GOLD |
| KAFFEE | GUMMI |
| KELLNER | KASPERL |
| KOCH | KLOSTER |
| KORKEN | KUGEL |
| KÜCHE | LINIE |
| LÖFFEL | MAGNET |
| MENÜ | MANTEL |
| MESSER | ORGEL |
| NUDEL | PAKT |
| OBER | PALAST |
| PFANNE | PFERD |
| PFEFFER | PILLE |
| PIZZA | Pinsel |
| RECHNUNG | RINDE |
| REZEPT | RING |
| SALAT | RUBIN |
| SALZ | RÜSSEL |
| SOSSE | SCHAUKEL |
| STUHL | SCHUH |
| SUPPE | SCHWAN |
| TABLETT | SPIEL |
| TASSE | SPINNER |
| TELLER | VEILCHEN |
| TISCH | WANNE |
| TOPF | WÜRFEL |
| TRINKGELD | ZIEGEL |
| WEIN | ZIRKUS |
